# EZH2 is essential for fate determination in the mammalian Isthmic area

**DOI:** 10.1101/442111

**Authors:** Iris Wever, Cindy M.R.J. Wagemans, Marten P. Smidt

## Abstract

The polycomb group proteins (PcGs) are a group of epigenetic factors associated with gene silencing. They are found in several families of multiprotein complexes, including Polycomb Repressive Complex (PRC) 2. EZH2, EED and SUZ12 form the core components of the PRC2 complex, which is responsible for the mono, di- and trimethylation of lysine 27 of histone 3 (H3K27Me3), the chromatin mark associated with gene silencing. Loss-of-function studies of *Ezh2*, the catalytic subunit of PRC2, have shown that PRC2 plays a role in regulating developmental transitions of neuronal progenitor cells; from self-renewal to differentiation and the neurogenic-to-gliogenic fate switch.

To further address the function of EZH2 and H3K27me3 during neuronal development we generated a conditional mutant in which *Ezh2* was removed in the mammalian isthmic (mid-hindbrain) region from E10.5 onward. Loss of *Ezh2* changed the molecular coding of the anterior ventral hindbrain leading to a fate switch and the appearance of ectopic dopaminergic neurons. The correct specification of the isthmic region is dependent on the signaling factors produced by the Isthmic organizer (IsO), located at the border of the mid- and hindbrain. We propose that the change of cellular fate is a result of the presence of *Otx2* in the hindbrain of *Ezh2* conditional knock-outs and a dysfunctional IsO, as represented by the loss of *Fgf8* and *Wnt1.* Our work implies that next to controlling developmental transitions, EZH2 mediated gene silencing is important for specification of the isthmic region by influencing IsO functioning and repressing *Otx2* in the hindbrain.

## Introduction

In recent years it has become apparent that next to transcription factors, gene regulation via epigenetics also plays an important part in the development of a multicellular organism (Bernstein et al., 2006; Mikkelsen et al., 2007; Mohn et al., 2008). Epigenetics can be defined as the biological processes that alter transcriptional activity by influencing the accessibility of the DNA by reorganizing the chromatin structure. Adaptation of chromosomal regions can rely on three distinct processes: (1) Modification of histones; (2) DNA methylation; and(3) non-coding RNAs mediated changes (Heesbeen et al., 2013). Histones contain numerous sites for different types of modifications, such as methylation, acetylation and phosporylation (Podobinska et al., 2017), which are highly dynamic and cause readily reversible changes in chromosomal organization (Cedar and Bergman, 2009). Methylation of histones can have different outcomes, while some sites correlate to transcriptional activity, others are associated with gene silencing (Podobinska et al., 2017). The polycomb repressive complex 2 (PRC2) mediates the mono, di and tri-methylation of lysine 27 of histone 3 (H3K27me1, 2 and 3) (Shen et al., 2008) and consists of three core subunits; Enhancer of zeste homolog 2 (EZH2), Embryonic ectoderm development (EED) and suppressor of zeste 12 (SUZ12), that are all crucial for the catalytic activity of the complex (Corley and Kroll, 2015). H3K27me3 is associated with gene silencing and shows a highly dynamic profile during development (Mohn et al., 2008). Together with histone 3 lysine 4 tri-methylation (H3K4me3), H3K27me3 is found on bivalent domains that silence developmental genes during one stage of development, but keeps them poised for activation in later stages of development (Bernstein et al., 2006; Mohn et al., 2008). The importance of H3K27 methylation and the PRC2 complex during development were shown in previous loss-of-function studies. The genetic ablation of one of the core subunits of the PRC2 complex led to a global loss of H3K27 di- and tri-methylation and a significant reduction in mono-methylation (Ferrari et al., 2014). In addition, knock-out (KO) embryos for one of the three core subunits of PRC2 fail to complete gastrulation due to defects in morphogenetic movements (Cao and Zhang, 2004a; Faust et al., 1998; O’Carroll et al., 2001). In later stages of development it was shown that when *Ezh2* was conditionally removed before the onset of neurogenesis in cortical progenitors the balance between self-renewal and differentiation was shifted toward differentiation (Pereira et al., 2010). In addition, neurogenesis was accelerated and onset of gliogenesis was earlier (Pereira et al., 2010). However, when *Ezh2* or *Eed* were removed during neurogenesis, the neurogenic phase was prolonged at the expense of the onset of astrogenesis (Hirabayashi et al., 2009). These data suggest that PRC2 plays a role in regulating developmental transitions in cortical progenitor cells; from self-renewal to differentiation and the neurogenic-to-gliogenic switch (Hirabayashi et al., 2009; Pereira et al., 2010). A similar role for *Ezh2* was found in neuronal progenitors (NPs) of the dorsal midbrain. *Wnt1Cre* driven deletion of *Ezh2* led to reduced number of NPs in the dorsal midbrain, due to elevated cell cycle exit and differentiation (Zemke et al., 2015). In addition to its role in regulating the transition from proliferation to differentiation, *Ezh2* was also found to be required for the maintenance of regional identity by suppressing forebrain traits (Zemke et al., 2015). To further address the function of EZH2 and H3K27me3 during neuronal development we generated a conditional mutant in which *Ezh2* was removed in the isthmic (mid-hindbrain) region from E10.5 onward (Sunmonu et al., 2009; Zemke et al., 2015). The mid-hindbrain region gives rise to several essential neurotransmitter systems, including dopaminergic (DA) and serotonergic (5-HT) neurons (Brodski et al., 2003). A critical event in the development of the different neuronal populations found in the midhindbrain region is the formation of the Isthmic Organizer (IsO) (Brodski et al., 2003; Crossley et al., 1996). The IsO secretes inductive signals that specify the mid-hindbrain region and it has been shown that a disorganized IsO influences the location and size of the DA and 5-HT system (Brodski et al., 2003; Kouwenhoven et al., 2016). The functioning and maintenance of the IsO is dependent on several feedback loops formed by multiple developmental factors, including *Fgf8*, *Wnt1, Lmx1b* and *En1* (Adams et al., 2000; Canning et al., 2007; Danielian and McMahon, 1996; Guo et al., 2006; Kouwenhoven et al., 2016; Martinez et al., 1999). In this study we show that the *En1cre* driven deletion of *Ezh2* and a consequential loss of H3K27me3 leads to a disorganized IsO and ectopic expression of the transcription factor *Otx2* in the hindbrain. In accordance with other studies in which *Otx2* expression is expanded caudally, DA cells are formed in rhombomere (R) 1 at the expense of R1 born neuronal subtypes, including 5-HT neurons (Brodski et al., 2003; Kouwenhoven et al., 2016; Sherf et al., 2015). Together, our data suggest that next to transcription factors, gene repression via EZH2 and H3K27me3 are required for the repression of Otx2 in the hindbrain, maintenance of the IsO and the correct fate-determination of the isthmic area.

## Material and Methods

### Ethics Statement

All animal studies are performed in accordance with local animal welfare regulations, as this project has been approved by the animal experimental committee (Dier ethische commissie, Universiteit van Amsterdam; DEC-UvA), and international guidelines.

### Animals

All lines were maintained on a C57BL/6J background (Charles River). *Ezh2*-floxed animals were generated by S.H. Orkin and a kind gift from F. Zilbermann (Friedrich Miescher Institute, Switserland) and have been previously described (Shen et al., 2008). *En1cre* animals were generated by A.L Joyner and a kind gift from S. Blaess (Rheinische Friedrich-Wilhelms-Universität, Gemany). *En1cre/+* animals were crossed with *En1cre-ERT +/+; R26RYFP/R26RYFP* to obtain *En1cre/+; R26RYFP/R26RYFP* (Kimmel et al., 2000). *Ezh2* L/L animals were crossed with *En1cre/+* or *En1cre/+; R26RYFP/R26RYFP* animals to obtain *En1cre/+; Ezh2* L/+ or *En1cre/+; Ezh2 L/+; R26RYFP/R26RYFP* animals. For the generation of embryos we crossed *En1cre/+; Ezh2 L/+* animals with *Ezh2 L*/+ animals and *En1cre/+; Ezh2 L/+ with En1cre/+; Ezh2 L*/+ animals for the generation of *En1/Ezh2* double mutant embryos. Embryos were isolated at embryonic day (E) 12.5, E14.5, considering the morning of plug formation as E0.5. Pregnant dams were euthanized by CO2 asphyxiation and embryos were collected in 1xPBS and immediately frozen on dry-ice (fresh frozen) or fixed by immersion in 4% paraformaldehyde (PFA) for 4-5hrs at 4°C. After PFA incubation, samples were cryoprotected O/N in 30% sucrose at 4°C. Embryos were frozen on dry-ice and stored at −80°C. Cryosections were slices at 16μm, mounted at Superfrost plus slides, air-dried and stored at −80°C until further use.

E14.5 *WnlCre/+; Ezh2 L/L* and wildtype littermate embryonic tissue were a kind gift from Prof. Dr. Filippo Rijli of the Friedrich Miescher Institute in Basel, Switzerland.

### Genotyping

The genotyping for the *Ezh2-flox* allele was executed with 50-100 ng of genomic DNA together with forward primer 5’- ACCATGTGCTGAAACCAACAG -3’ and reverse primer 5’- TGACATGGGCCTCATAGTGAC – 3’ resulting in a 395 bp product for the wild type allele and a 361 bp product for the *floxed* allele.

Genotyping of the *En1cre* allele was performed with 50-100 ng of genomic DNA together with primer pair En1Cre 5UTR_F3 5’-CTTCGCTGAGGCTTCGCTTT-3’ and En1Cre Cre_R2 5’-AGTTTTTACTGCCAGACCGC-3’ resulting in a product at 240 bp for the *Cre*-allele.

Genotyping for the *R26R-YFP* allele was performed using 3 primers Rosa_mutant primer 5’-AAGACCGCGAAGAGTTTGTC -3’, Rosa_wildtype primer 5’- GGAGCGGGAGAAATGGATATG -3’ and a Rosa_common primer 5’- AAAGTCGCTCTGAGTTGTTAT–3’ with 50-100 ng of genomic DNA. The PCR reaction gave a product at 320 bp for the mutant *R26R-YFP* allele and a product of 600 bp for the wildtype allele.

### In situ hybridization

In situ hybridization with digoxigenin (DIG)-labeled RNA probes was performed as described (Smidt et al., 2004; Smits et al., 2003). The following DIG-Labeled RNA probes were used; *Th* (Grima et al., 1985), *Pitx3, Nurr1* (Smidt et al., 2000), *Ahd2* (Jacobs et al., 2007), *En1, Cck* (Jacobs et al., 2011), *Lmx1a* (Hoekstra et al., 2013), *Wnt1* (Mesman et al., 2014), *Fgf8, Otx2* (Kouwenhoven et al., 2016) and a *Nkx6.1* probe containing bp 1182-1713 (532bp) of the mouse cDNA sequence (NM_144955.1).

### Fluorescence Immunohistochemistry

Cryosections were blocked with 4% HIFCS in THZT buffer [50mM Tris-HCL, pH 7.6; 0.5M NaCl;0.5% Triton X-100] and incubated with a primary antibody; Th [Rabbit-TH (Pelfreeze 1:1000), Sheep-TH (Millipore AB1542, 1:750)], EYFP [Chicken-GFP (Abcam, 1:500)], H3K27me3 [Rabbit-anti-H3K27me3 (Millipore, 17-622 1:2000)] or Serotonin [Rabbit-anti-Serotonin (Immunostar, 1:500)], in THZT buffer overnight at room temperature. The following day the slides were washed and incubated for 2 hrs at room temperature with secondary Alexafluor antibody (anti-rabbit, anti-sheep and antichicken (In Vitrogen, 1:1000) in Tris buffer Saline (TBS). After immunostaining nuclei were stained with DAPI (1:3000), embedded in Fluorsave (Calbiogen) and analyzed with the use of a fluorescent microscope (Leica). All washes were done in TBS and double staining was performed sequential. The antibody against H3K27me3 required antigen retrieval, which was executed as follows; slides were incubates 10 minutes in PFA after which they were submersed in 0.1M citrate buffer pH 6.0 for 3 minutes at 1000Watts followed by 9 minutes at 200Watts. Slides were left to cool down, after which protocol was followed as described above.

### Quantitative PCR (qPCR)

RNA was isolated from dissected E12.5 or E14.5 midbrains of *En1cre/+; Ezh2 +/+* and *Ezh2* cKO embryos. RNA was isolated with Trizol (ThermoFisher) according to the manufacturers protocol. For E12.5 a single midbrain was used per sample (*En1cre/+; Ezh2 +/+* n=3 and *En1cre/+; Ezh2* L/L n=4) For E14.5 two midbrains were pooled for the *Ezh2* cKO samples and a single midbrain was used per sample for the wildtype (wildtype n=4 and *En1cre/+; Ezh2* L/L n=3). Relative expression levels were determined by using the QuantiTect SYBR green PCR lightCycler kit (Qiagen) according to the manufacturers instructions. For each reaction 10ng (dissected midbrain) of total RNA was used as input. Primers used for *En1* were previously published (Jacobs et al., 2011)., Further primer designs: *Fgf8:* forward 5’-GCAGAAGACGGAGACCCCT and reverse 5’-CGTTGCTCTTGGCAATTAGCTTCC (product size 136bp), *Wnt1* forward 5’-CAGCAACCACAGTCGT CAGA and reverse 5’-TTCACGATGCCCCACCATC (produce size 170bp) and *Otx2:* forward 5’-GGGAAGAGGTGGCACTGAAA and reverse 5’-CTTCTTGGCAGGCCTCACTT (produce size 137bp).

## Results

### *En1Cre* driven deletion *of Ezh2* causes a widespread loss of H3K27me3

As described above, EZH2 functions as the methyltransferase of PRC2, which catalyzes the methylation of H3K27 (Cao and Zhang, 2004b; Margueron et al., 2008; Shen et al., 2008). Early deletion of *Ezh2* has been shown to cause an overall ablation of H3K27me3 and transitions in developmental phases, like the switch from self-renewal to differentiation, are disrupted (Hirabayashi et al., 2009; Pereira et al., 2010; Zemke et al., 2015). In the mid-hindbrain region several neuronal substypes arise, including dopaminergic (DA) and serotonergic (5-HT) neurons (Brodski et al., 2003; Kouwenhoven et al., 2016). An essential event in the emergence of these neuronal populations is the formation of the IsO, which secretes inductive signals that determine the size and location of the different neuronal subtypes originating in the mid-hindbrain region (Brodski et al., 2003; Crossley et al., 1996; Kouwenhoven et al., 2016). In this study we aim to gain further insight into the role of EZH2 and PRC2 functioning in the development of the mid- hindbrain region by generating a conditional knock-out (cKO) mouse. *Ezh2* was removed from the mid- hindbrain region from E10.5 onward by crossing *Ezh2*-floxed mice (Shen et al., 2008) with *En1cre* animals (Kimmel et al., 2000). The CRE recombinant region, as visualized by YFP *(R26RYFP)* (Srinivas et al., 2001), extends rostrally into the ventral region of prosomere 3 and caudally to the presumed R1/R2 border (**Figure 1A**) (Kouwenhoven et al., 2016). To determine whether the deletion of *Ezh2* in the mid-hindbrain leads to the loss of H3K27me3 an immunohistochemistry for H3K27me3 was performed in combination with DAPI staining (**Figure 1B**). In *En1cre/+; Ezh2* L/L animals H3K27me3 could not be observed in the midhindbrain region from E12.5 onward (**Figure 1B** **(3)**, arrowheads), while the mark could be detected in this region in wildtype embryos (**Figure 1B** **(1)**). In addition, H3K27me3 is present in the *Ezh2* cKO outside of the CRE recombination region (**Figure 1B** **(4)**), indicating that the deletion of *Ezh2* is specific for the *En1* positive area and that the ablation of *Ezh2* alone is sufficient to remove H3K27me3 in this region.

**Figure 1:**
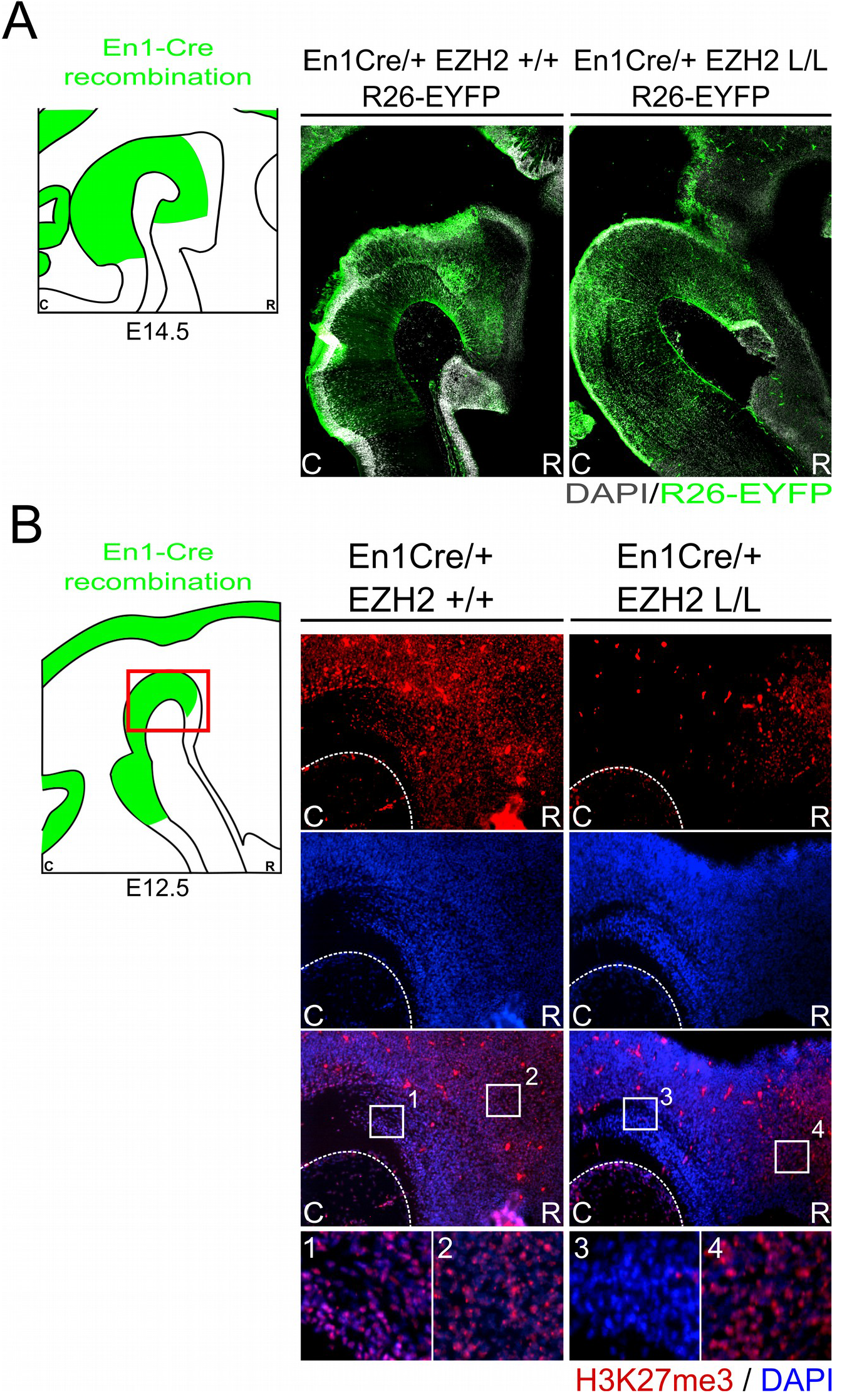
H3K27me3 is lost in the *En1* expression domain of *En1Cre/+; Ezh2* L/L E12.5 Embryos. (A) Immunohistochemistry of R26R-YFP (green) was performed in E14.5 wildtypes and *Ezh2* cKO embryos to determine the CRE recombinant area. R26R-YFP expression is found from the ventral region of P3 and extended caudally up until the R1/R2 border. (B) The presence of H3K27me3 was determined in E12.5 *En1cre/+; Ezh2 +/+* and *En1cre/+; Ezh2* L/L embryos by means of immunohistochemistry. H3K27me3 (red) was lost in the midbrain of *Ezh2* cKO embryos (3), but was still present in the dorsal pretectum (4). Nuclear localization of H3K27me3 was verified by colocalization with DAPI staining (blue).

### Ectopic dopaminergic neurons arise in the hindbrain of *En1Cre/Ezh2* L/L animals

One of the essential neuronal subtypes that develops in the Isthmic region are mesodiencephalic dopaminergic (mdDA) neurons. MdDA progenitors originate from the floor plate of the ventral midbrain under the influence of intersecting signals of FGF8, OTX2 and WNT1 (Mesman et al., 2014; Ono et al., 2007; Placzek and Briscoe, 2005). During neurogenesis, mdDA progenitors enter the G0 phase of the cells cycle to give rise to post-mitotic mdDA precursors that start migrating towards their final diencephalic (Prosomere 1-2) and midbrain domains (Kawano et al., 1995; Mesman et al., 2014; Shults et al., 1990). During this phase mdDA precursors differentiate further into mdDA neurons characterized by the presence of tyrosine hydroxylase (TH), the rate-limiting enzyme in dopamine synthesis. Although most TH+ cells are born around E11.0 (Bayer et al., 1995), it is not until E14.5 that they express most genes that define a mature mdDA neuron, like *Dat, Ahd2* and *Cck (Arenas et al., 2015; Iversen, 2010; Veenvliet et al., 2013)*. To examine whether the loss of *Ezh2* in the mid- hindbrain region affects the DA neuronal population we performed immunohistochemistry for TH at E14.5 (**Figure 2**). When comparing *En1cre/+; Ezh2 +/+* embryos to *En1cre/+; Ezh2* L/L embryos a caudal expansion of TH+ neurons is observed (**Figure 2**, arrowheads). The biggest changes in the TH+ domain are observed in the medial sections where a group of TH+ neurons is found in an ectopic location caudal to the midbrain.

**Figure 2:**
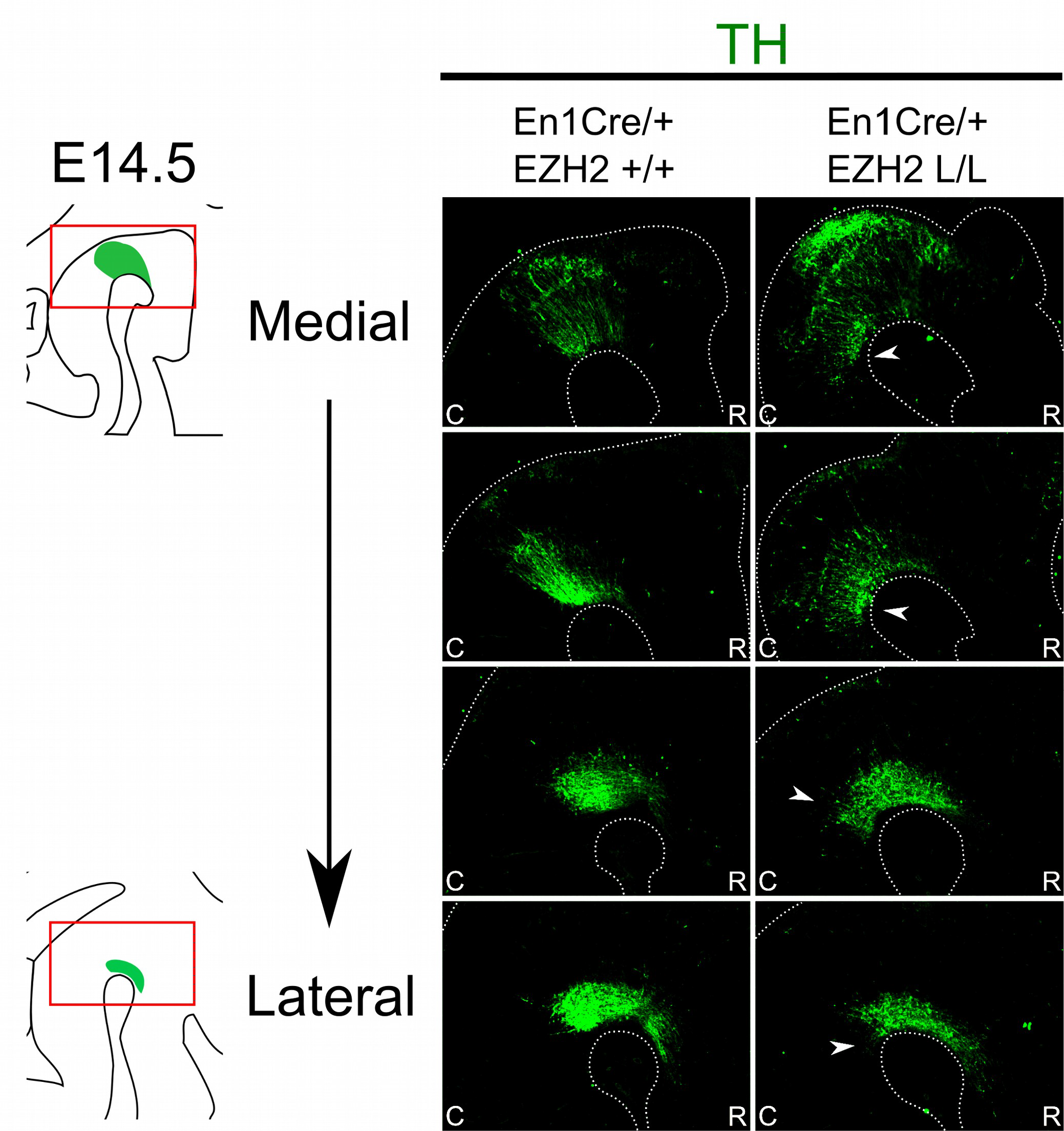
*En1cre* driven deletion of *Ezh2* leads to a caudal expansion of the TH+ domain. Protein expression of TH (green) was evaluated by means of immunohistochemistry. In E14.5 *En1cre/+; Ezh2* L/L embryos the TH+ domain is shifted caudally in comparison to *En1Cre/+; Ezh2 +/+* embryos (arrowheads). The caudal expansion of the TH+ domain is most apparent in medial sections (upper panels).

Next to mdDA neurons, the mid-hindbrain region also gives rise to 5-HT neurons (Alonso et al., 2013; Brodski et al., 2003; Fox and Deneris, 2012; Sherf et al., 2015). 5-HT neurons develop caudally to the IsO and are arranged into three distinct nuclei; the dorsal raphe nucleus (DRN), the median raphe nucleus (MRN) and the prepontine raphe nucleus (PpnR), which are R2-derived (Alonso et al., 2013; Fox and Deneris, 2012). Analysis of serotonin by immunohistochemistry at E14.5 showed that the cytoarchitecture of the 5-HT system is changed in *Ezh2* cKOs (**Figure 3A**). Under normal conditions, no 5-HT+ cells can be found in between the DRN and the MRN where the decussation of the superior cerebellar peduncle runs, however in *En1cre/+; Ezh2 L/L* embryos 5-HT+ neurons are present in this region (**Figure 3A**, arrowheads). In addition to the disorganized structure, a loss of cells in the DRN is observed in lateral sections of *En1cre/+; Ezh2* L/L embryos when compared to wildtypes (**Figure 3A**, arrows). Previous studies have already shown that the caudal expansion of the DA domain negatively affects the generation of 5-HT+ neurons in R1 (Brodski et al., 2003; Kouwenhoven et al., 2016; Sherf et al., 2015). So to get a better insight into the effect of the caudal expansion of the TH+ domain into the ventral hindbrain on the generation of the 5-HT+ population in this region, we performed a double immunohistochemistry for 5-HT and TH on adjacent slides (**Figure 3B**). In line with these studies, no 5-HT+ cells were found dorsally of the ventral ectopic patch of TH+ neurons or directly caudal to the dorsal DA domain in the *Ezh2* cKOs (**Figure 3B** **(2)**, arrowhead). In wildtypes the 5-HT+ population of the DRN align the dorsal DA domain (**Figure 3B** **(1)**, arrowhead) and the lack of cells dorsal of the ectoptic TH+ patch suggests that the caudal expansion of the TH+ domain is at the expense of the 5-HT+ domain corresponding to the DRN.

**Figure 3:**
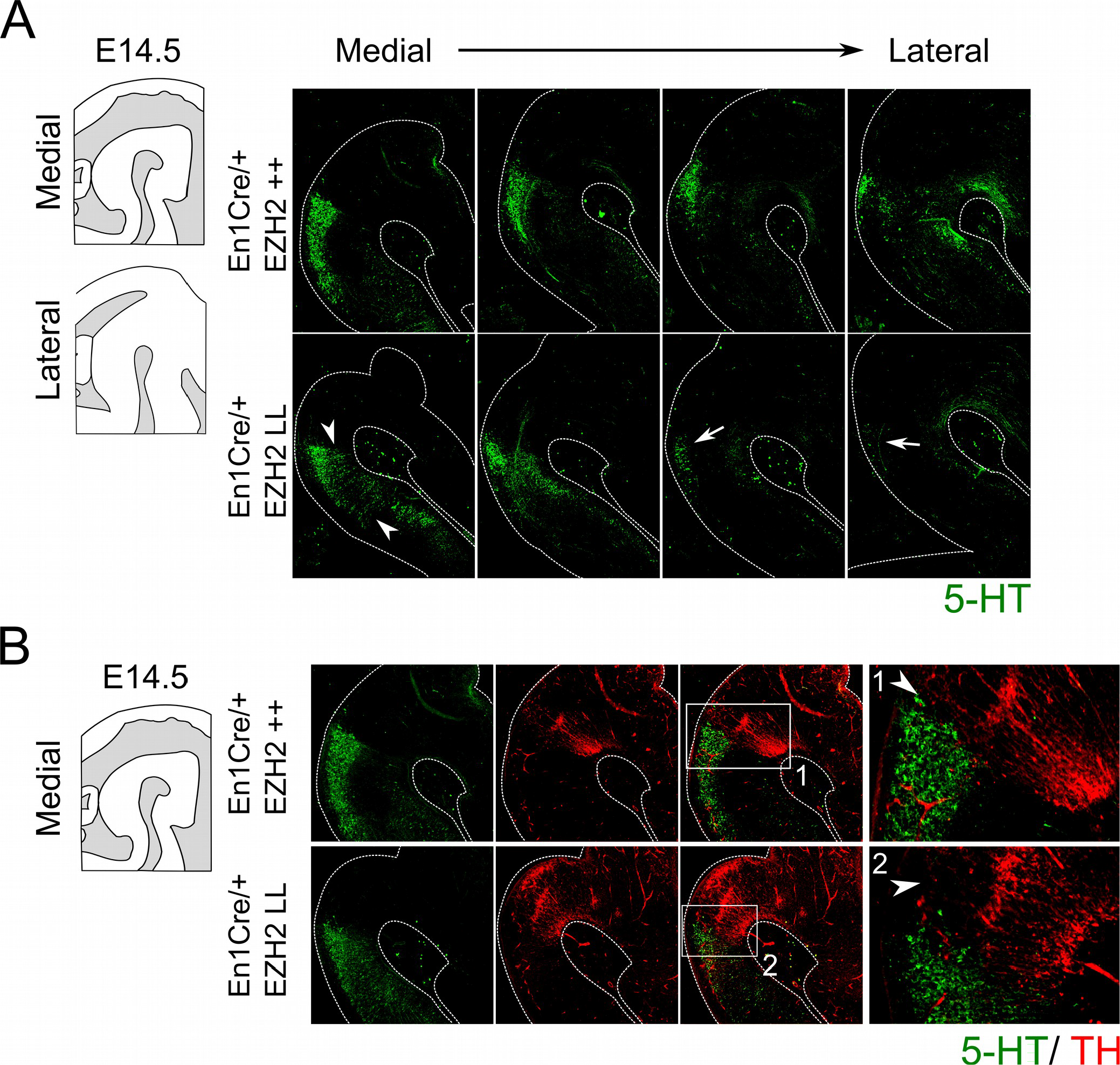
Genetic ablation of *Ezh2* affect the cytoarchitecture of the 5-HT population and leads to the loss of neurons of the Dorsal Raphe Nucleus. (A) The 5-HT positive population was examined by performing a immunohistochemistry for 5-HT (green). The overall cytoarchitecture of the 5-HT population is affected (arrowheads) and cells of the DRN are lost in the lateral sections of *En1cre/+; Ezh2* L/L E14.5 embryos (arrows). (B) A double immunohistochemistry for 5-HT (green) and TH+ (red). (1) In wildtypes 5-HT cells align the dorsal mdDA domain (arrowhead), while in *En1cre* driven *Ezh2* cKOs (2) TH+ cells are found in the hindbrain and the 5-HT cells of the DRN cells are lost dorsal of the ectopic TH+ patch (arrowhead).

Together our data indicate that *En1cre* driven deletion of *Ezh2* leads to the appearance of TH positive cells caudal to the midbrain and that this extension of the TH+ domain is at the expense of 5-HT+ neurons.

### Conditional deletion of *Ezh2* changes the molecular coding of the anterior hindbrain

To further substantiate whether the ectopic TH+ cells are DA-like neurons we performed *In situ* hybridization for several DA marks (**Figure 4**). *Lmx1a, Nurr1 and Pitx3* have been shown to play an important role in mdDA development (Deng et al., 2011; Jankovic et al., 2005; Saucedo-Cardenas et al., 1998; Yan et al., 2011). While *Lmx1a* has been shown to be required for the activation of neurogenesis and for the suppression of alternative cells fates (Andersson et al., 2006; Omodei et al., 2008), *Nurr1* induces the DA neurotransmitter phenotype (Saucedo-Cardenas et al., 1998; Smits et al., 2003). *Pitx3* expression is initiated in later stages of development and has been shown to be important for the specification of the different mdDA subsets and for the survival of the DA neurons of the Substantia Nigra pars compacta (SNc) (Hwang et al., 2003; Maxwell et al., 2005; Smidt et al., 1997; Veenvliet et al., 2013). The spatial expression of *Th* shows a corresponding expression pattern as was observed with immunohistochemistry, an ectopic patch of expression caudal to the midbrain (arrowhead). Analysis of *Lmx1a* and *Nurr1* expression on adjacent sections demonstrated that they are also expressed in the ectopic *Th* area caudal to the midbrain in *Ezh2* cKO embryos (**Figure 4A (*)**, arrowheads). In addition, *Pitx3* was also found ectopically, caudal to the midbrain in *En1cre/+; Ezh2* L/L embryos (**Figure 4A (*)**, lowest panel, arrowhead), indicating that the ectopic TH+ cells have a DA-like phenotype. Next to DA marks that are expressed by all mdDA neurons we also examined the expression pattern of two subset marks, *Cck* and *Ahd2* (**Figure 4B**) (Veenvliet et al., 2013). *Cck* is normally expressed by the caudomedial mdDA neuronal population and when examining *En1cre/+; Ezh2 +/+* and *Ezh2* cKO midbrains, expression was also detected caudal to the midbrain in *Ezh2* cKO embryos (**Figure 4B**, left panel, arrowhead). Interestingly, *Adh2* expression was also sparsely observed caudal to the midbrain in medial sections of *En1cre/+; Ezh2* L/L embryos (**Figure 4B**, right panel arrowhead), while *Ahd2* is restricted to the rostrolateral DA population in wildtypes (Veenvliet et al., 2013).

**Figure 4:**
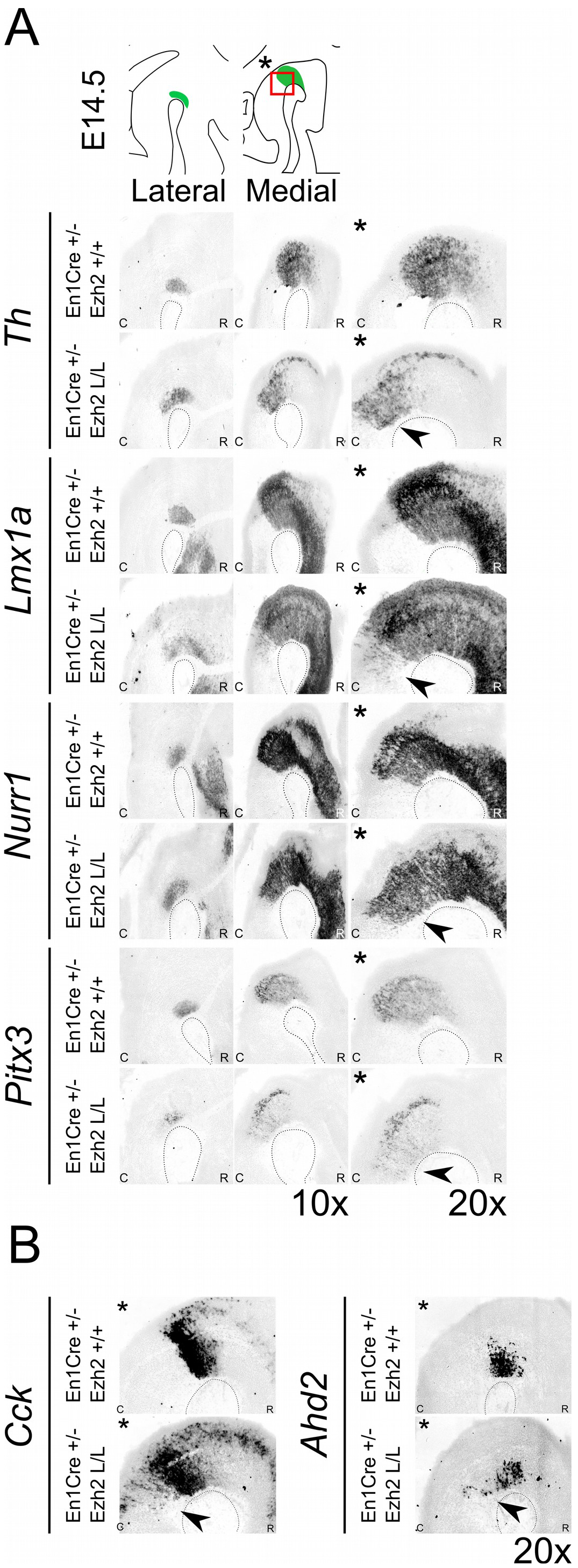
DA marks can be detected more caudally in *Ezh2* cKO animals. Analysis of DA marks, *Th, Lmx1a, Nurr1, Pitx3* and subset marks *Cck* and *Ahd2* at E14.5 by means of *in situ* hybridization (A, B).(A) An ectopic patch expressing *Th, Lmx1a, Nurr1* and *Pitx3* can be observed caudal to the midbrain in the *Ezh2* mutant, which is not present in *En1cre/+; Ezh2* +/+ animals (arrowheads). (B) In addition to general marks the caudomedial subset mark *Cck* and the rostrolateral mark *Ahd2* were also found to be shifted caudally in *En1cre/+; Ezh2 L/L* animals compared to wildtypes (arrowheads).

In normal conditions, *Nkx6.1* expressing trochlear motor neurons are located in and directly caudal to the IsO (Chandrasekhar, 2004; Prakash et al., 2009). As described above, *Lmx1a* has been demonstrated to suppress alternative cell fates, including motor neurons by the repression of *Nkx6.1* (Andersson et al., 2006). The presence of *Lmx1a* and other DA related genes in ventral R1 suggest that the molecular coding in this region might be altered, leading to a DA-like cell fate. To assess whether the deletion of *Ezh2* leads to an alternative cell fate and the suppression of *Nkx6.1*, we assessed the presence of *Nkx6.1* in the ventral Isthmic area by means of *in situ* hybridization (**Figure 5**). When examining the midhindbrain region of *En1cre/+; Ezh2* L/L embryos almost all *Nkx6.1* staining was lost in the Isthmic area (**Figure 5** **(2)**, arrowhead), corresponding to the region where normally the trochlear motor neurons develop (**Figure 5** (1)) (Chandrasekhar, 2004, 2004). In contrast, *Nkx6.1* expression in other regions, like the red nucleus, were similar to *En1cre/+; Ezh2* +/+ littermates, indicating that the loss of *Ezh2* specifically affects *Nkx6.1* expression in the anterior hindbrain.

**Figure 5:**
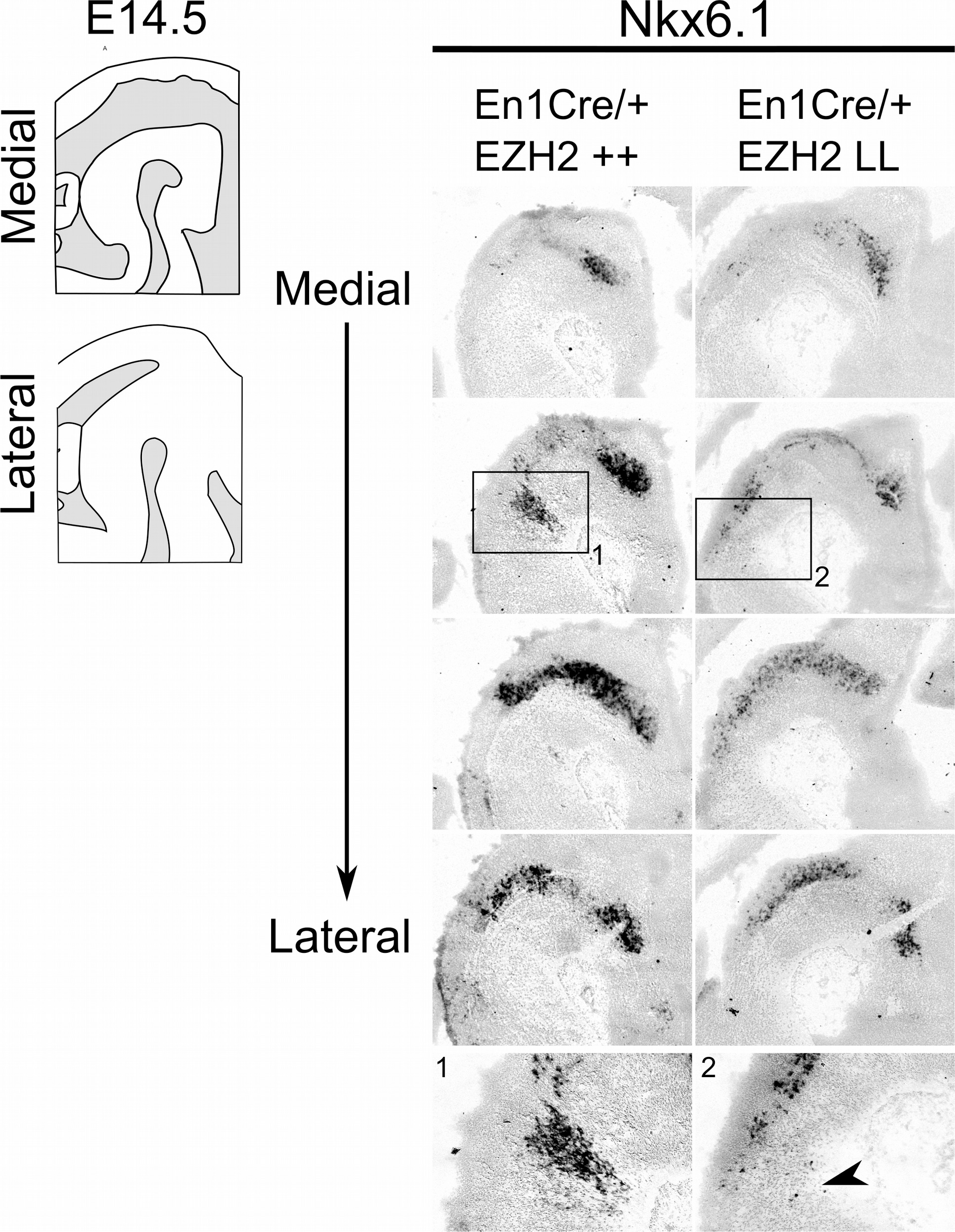
*Nkx6.1* expression is lost in the ventral R1 region in *Ezh2* cKO E14.5 embryos. *In situ* hybridization for *Nkx6.1* was performed on E14.5 *En1cre/+; Ezh2 +/+* and *Ezh2* cKO embryos. Analysis of the spatial expression of *Nkx6.1* shows that the expression is lost in the ventral region of R1 in *En1cre/+; Ezh2* L/L embryos (1 and 2, arrowhead), but not in the midbrain.

Together, these results suggest that the loss of *Ezh2* changes the molecular coding of the anterior hindbrain, leading to the presence of an ectopic patch of dopamine-like cells caudal to the midbrain and the loss of *Nkx6.1+* expression in this region.

### *En1Cre* driven *Ezh2* cKOs show a similar phenotype as *En1* mutants independent of *En1*

Previous studies have shown that in the absence of *En1*, DA-related genes display a caudal expansion of their expression domain in more medial sections. The presence of these ectopic DA cells can be detected as early as E12.5 and they show a similar molecular and electrophysiological profile as mdDA neurons (Kouwenhoven et al., 2016; Veenvliet et al., 2013). The *Ezh2* cKO is heterozygous for *En1* and to verify that the ectopic presence of DA neurons caudal to the midbrain in the *En1cre/+; Ezh2* L/L is not a consequence of the combined loss of *En1* and *Ezh2* we verified the levels and spatial expression of *En1* via qPCR and *In situ* hybridization at E14.5 in *En1cre/+; Ezh2* L/L embryos (**Supplemental Figure 1A, B**) and performed a second set of experiments on *Wnt1Cre* driven *Ezh2* cKOs (**Figure 6**) (Kratochwil et al., 2017). When examining the spatial expression of *En1* in *En1cre/+; Ezh2* L/L embryos a similar pattern of expression as in *En1cre/+; Ezh2 +/+* animals is observed (**Supplemental Figure 1A**). However, the data also suggest that in the medial sections, *En1* is ectopically expressed in the ventral R1 region of *En1cre* driven *Ezh2* cKOs (**Supplemental Figure 1A**, arrowhead), as was found for Nurr1, Lmx1a and Pitx3 as shown above. Quantitative PCR analysis indicated that the level of expression was not significantly changed between *En1cre/+; Ezh2 +/+* and *En1cre/+; Ezh2* L/L animals (**Supplemental figure 1B**). *Wnt1Cre/+; Ezh2* L/L embryos have been studied before and a previous study on the developing midbrain of these animals showed that *Wnt1Cre* driven deletion of *Ezh2* leads to a loss of H3K27me3 in the Cre-recombinant area from E12.5 onward (Zemke et al., 2015). In addition, they showed full *Wnt1Cre*-mediated recombination in the caudal midbrain and R1 and while they used the *Wnt1Cre; Ezh2* model to study early development of the dorsal midbrain, the presence of CRE-recombinase in the caudal midbrain and R1 makes it possible for us to also use this model to validate our initial findings observed in *En1cre* driven *Ezh2* cKOs independent of the dosage of En1. (Zemke et al., 2015). We first performed immunohistochemistry for TH at E14.5 (**Figure 6A**). Similar to the *En1cre/+; Ezh2* L/L phenotype, we observed a group of TH+ cells caudal to the midbrain (**Figure 6A**, arrowheads). In addition to *Th* expression we also verified the presence of several other DA marks by means of *In situ* hybridization (**Figure 6B, C**). Next to TH protein, *Th*, *Pitx3* and *Cck* were found to be expanded caudally in the medial sections of the *Wnt1Cre; Ezh2* L/L compared to wildtypes (**Figure 6B, C**, arrowheads). However, in contrast to *En1cre; Ezh2* L/L animals almost no *Ahd2* expression was detected in the *Wnt1Cre; Ezh2* L/L embryos in comparison to the *Wnt1Cre/+; Ezh2 +/+* embryos (**Figure 6C**, lower panel, arrows).

**Figure 6:**
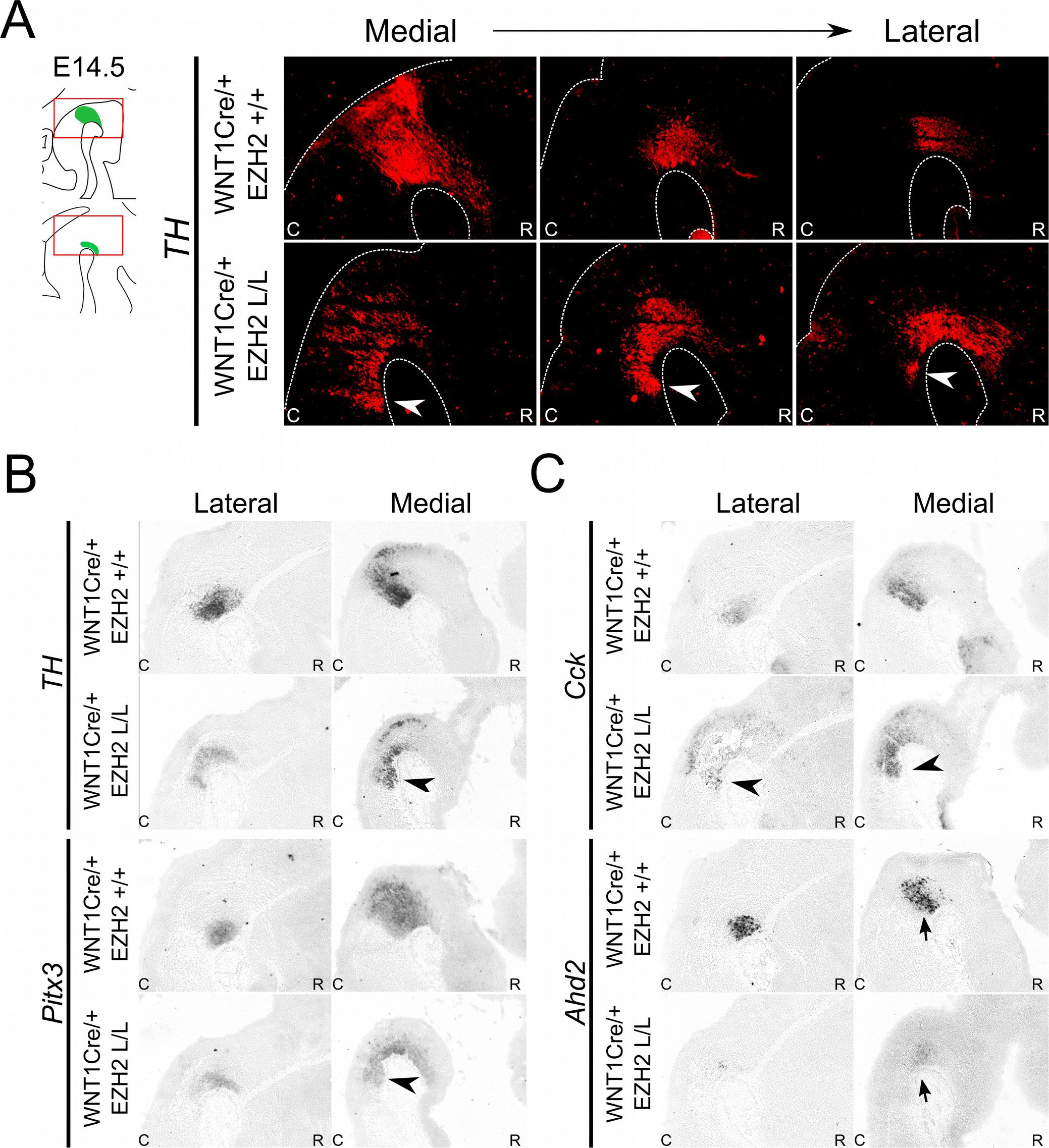
*Wnt1Cre* driven deletion of *Ezh2* results in a comparable presence of ectopic DA cells as observed in the *En1Cre/+; Ezh2* L/L. (A) Immunohistochemistry for TH (red) demonstrates that TH+ cells are present caudal to the midbrain in *Wnt1Cre/+; Ezh2* L/L embryos at E14.5. The ectopic TH+ neurons can be observed in lateral and medial sections (arrowheads). (B) Analysis of spatial expression of *Th, Pitx3, Cck* and *Ahd2* in adjacent E14.5 sections by means of *in situ* hybridization. Expression of *Th, Pitx3* and *Cck* is found in a similar location caudal to the midbrain (arrowheads). Expression of *Ahd2* is reduced the both the lateral and medial sections (arrows).

To further investigate a possible relationship between the phenotypes of *En1* ablation (Kouwenhoven et al., 2016) and *Ezh2* ablation we investigated whether genetic removal of *En1* influences the presence of H3K27me3 in the mid- hindbrain region. Therefore, an immunohistochemistry experiment was performed in *En1* mutant animals (**Figure 7A**). H3K27me3 was found in the mid-hindbrain region of both the *En1* mutants and wildtype controls. In addition, H3K27me3 was also detected in the ectopic DA cells in the *En1* mutant (**Figure 7A (*)**, arrowheads), suggesting that *Ezh2* and general PRC2 activity is not affected in these cells. Next to analyzing the individual mutants, we also generated double mutants by crossing *En1cre/+; Ezh2* L/+ with *En1cre/+; Ezh2* L/+. Immunohistochemistry on TH was performed to analyze the DA population in the resulting mutants (**Figure 7B**). In *En1creCre; EZh2* L/L embryos the DA domain expanded even further into the hindbrain than in the *En1cre/+; Ezh2* L/L embryos (**Figure 7B**, arrowheads) and the overall cytoarchitecture is more disorganized in the double mutant when compared to both the *Ezh2* cKO and the wildtype (**Figure 7B**, arrows), suggesting that *En1* and *Ezh2* do have a functional convergence although the molecular mechanism towards that convergence is not identical.

**Figure 7:**
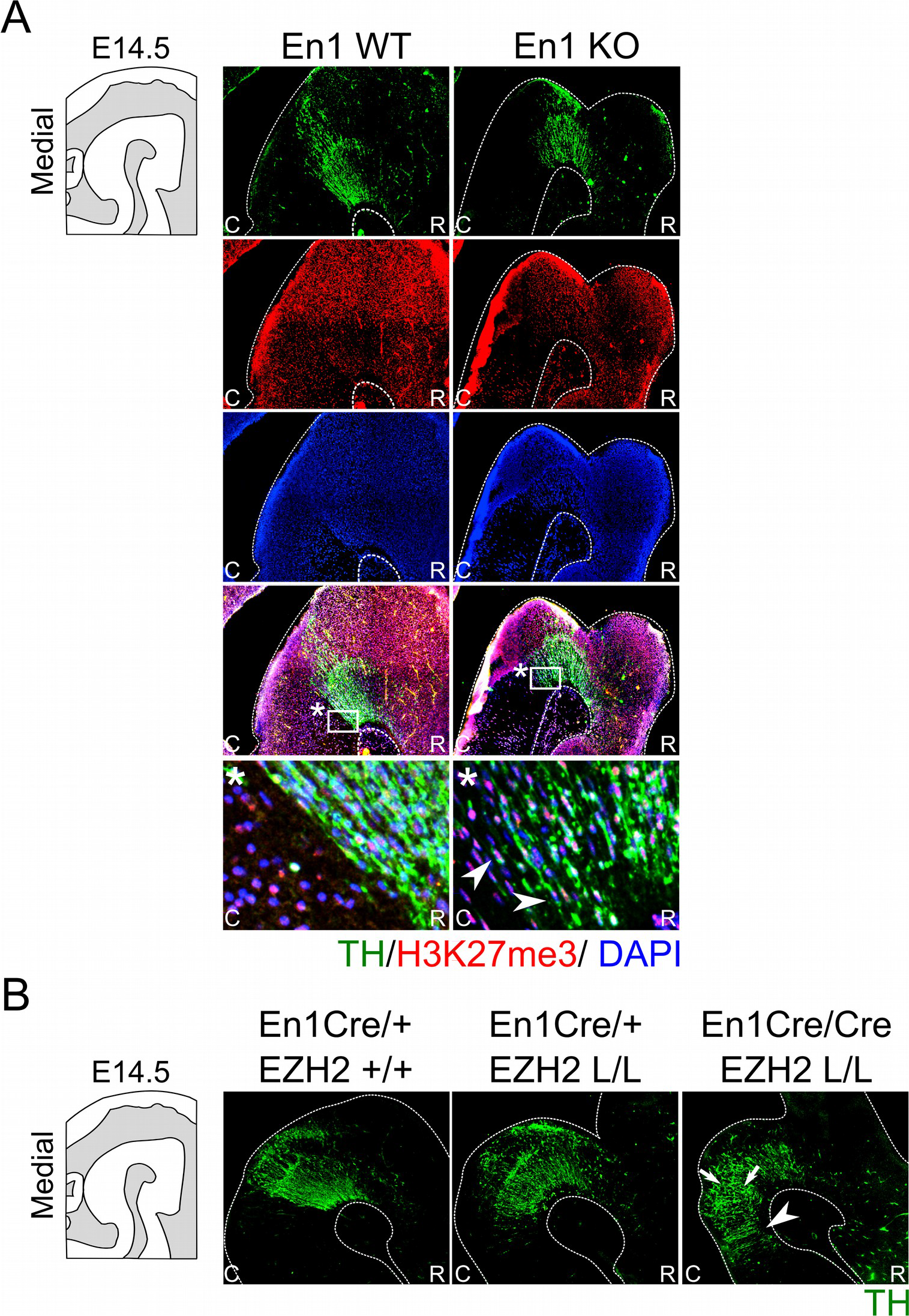
H3K27me3 is present in En1-deficient ectopic DA cells, whereas *En1/Ezh2* double mutants display an aggravated caudal expansion of the DA domain. The presence of H3K27me3 in ectopic DA cells observed in En1 mutants was investigated by means of a double immunohistochemistry for TH (green) and H3K27me3 (red) in wildtypes and *En1* −/− littermates. At E14.5 the TH+ cells found ectopically in the hindbrain of *En1* mutant are also positive for H3K27me3 (arrowheads). Nuclear localization was confirmed by performing a DAPI staining (blue). (B) Immunohistochemistry for TH (green) was performed on wildtypes, *Ezh2* cKOs and *En1creCre; Ezh2* L/L double mutant animals. TH+ cells can be found further caudal into the hindbrain in double mutants compared to both the *En1Cre/+; Ezh2* L/L and *En1Cre/+ Ezh2 +/+* animals (arrowheads). In addition the overall cytoarchitecture is disorganized in *En1creCre; Ezh2* L/L animals(arrows).

In summary, our data demonstrates that the loss of *Ezh2* and H3K27me3 in the mid-hindbrain region lead to a caudal expansion independently of *En1* expression. In addition, we show that although *En1* mutant animals have a similar phenotype, global H3K27me3 is not affected in these mutants, suggesting that overall EZH2 functioning is not affected in *En1* mutants in the isthmic area.

### *Ezh2* influences IsO determinants

The IsO is located at the border between the mid-and hindbrain and has been shown to be critical for the specification of both the midbrain as the anterior hindbrain (Brodski et al., 2003; Crossley et al., 1996; Joyner et al., 2000). The location of the IsO is initially determined by the mutual repression of two opposing factors, *Otx2* and *Gbx2* (Brodski et al., 2003; Wurst and Bally-Cuif, 2001). Loss- and gain-of-function showed that when the IsO is moved over the anterior-posterior axis the size of the two main neurotransmitter systems formed in this region, dopaminergic and serotonergic, is affected. When the IsO is moved to a more anterior position, the serotonergic system expands into the midbrain, while a caudal shift causes an expansion of the DA system into the hindbrain (Brodski et al., 2003; Kouwenhoven et al., 2016). A disorganization of the IsO might explain the caudal shift of the DA population observed in *En1cre/+; Ezh2* L/L embryos. We performed *in situ* hybridization of several genes associated with IsO functioning to determine whether the IsO was affected in *Ezh2* cKOs (**Figure 8**). *Fgf8* mediates the inducing capacity of the IsO (Crossley et al., 1996; Joyner et al., 2000; Martinez et al., 1999) and it expression can be observed as a transverse ring that encircles the neural tube in the vicinity of the mid-hindbrain border (**Figure 8A**, upper panel) (Martinez et al., 1999).

**Figure 8:**
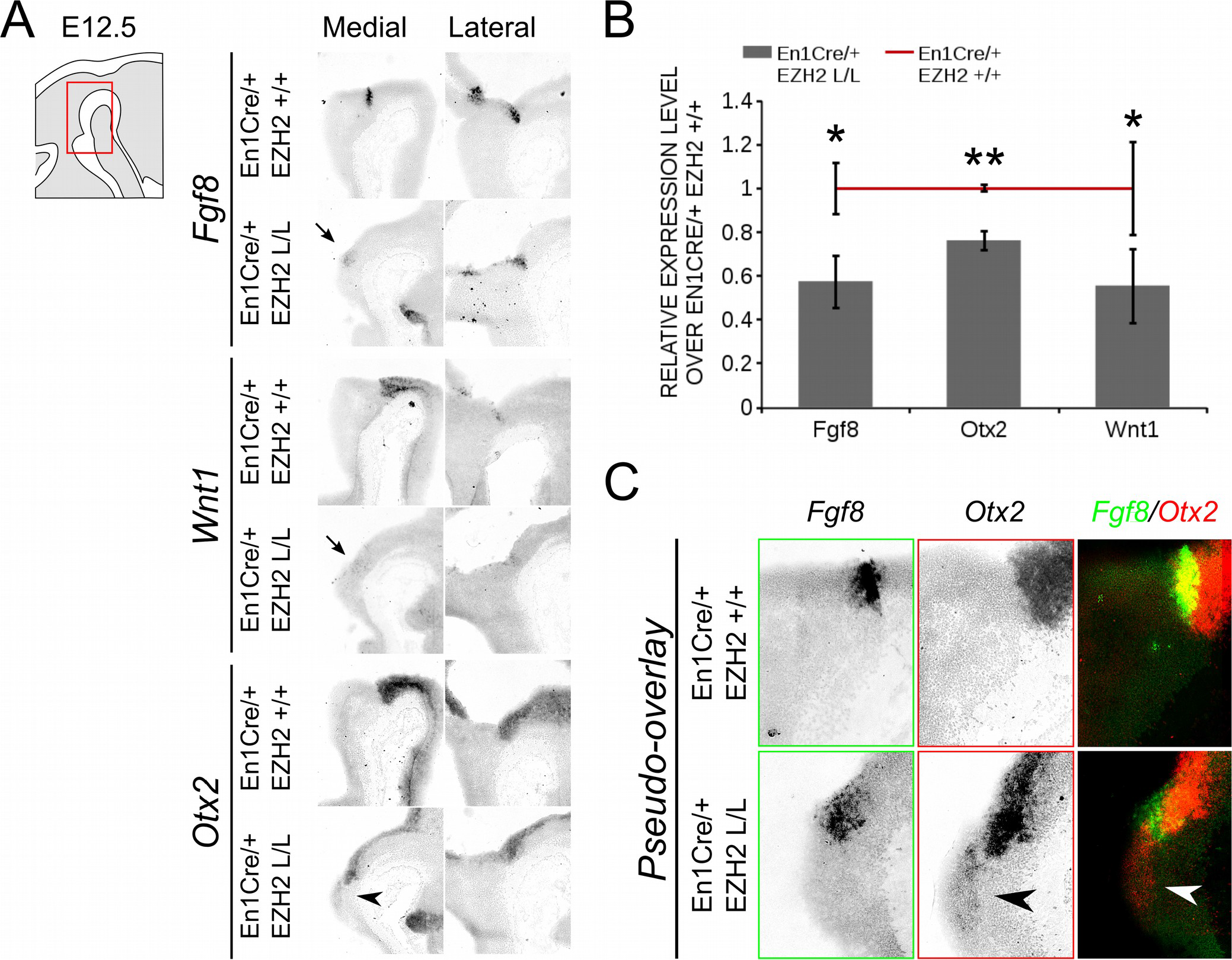
*Otx2* expression can be found caudal to the IsO in the hindbrain of *En1Cre/+; Ezh2* L/L E12.5 embryos. (A) The expression of *Fgf8, Otx2* and *Wnt1* was analyzed in E12.5 wildtypes and *Ezh2* cKO embryos via *in situ* hybridization. *Otx2* expression was found in the hindbrain of medial sections (arrowhead)m while expression of both *Fgf8* and *Wnt1* was reduced in the medial sections of E12.5 *En1cre/+; Ezh2* L/L embryos (arrows). (B) qPCR for *Fgf8, Otx2* and *Wnt1* comparing the wildtype (n=3, red line) expression levels to the *En1cre/; Ezh2* L/L (n=4, grey bars) levels. All factors were significantly down-regulated in the *Ezh2* cKO, *Fgf8* is ~ 43% reduced (* P<0.05, one-tailed), *Otx2* levels were reduced to ~ 76% (** P< 0.01, one-tailed) and ~ 45% of the *Wnt1* expression is lost (* P< 0.05, one-tailed). (C) Pseudo-overlay of adjacent slides for *Fgf8* and *Otx2* demonstrates that *Otx2* is found caudal of *Fgf8* expression in *Ezh2* cKO animals but not in controls.

When comparing *En1cre/+; Ezh2 +/+* to *En1cre/+; Ezh2* L/L mid-hindbrain sections, we observed a partial loss of expression in medial sections of E12.5 *En1cre/+; Ezh2* L/L embryos (**Figure 8A**, upper panel, arrow), while lateral no clear changes in the expression pattern could be observed. The reduction of the levels of *Fgf8* were verified using qPCR (n=3, P<0.05, one-tailed) (**Figure 8B**). In addition to *Fgf8*, we also studied the expression pattern of *Wnt1. Wnt1* is essential for the maintenance of several genes associated with IsO functioning, including *Fgf8* and *En1* (Canning et al., 2007; Danielian and McMahon, 1996; McMahon et al., 1992). Similar to *Fgf8*, reduced expression of *Wnt1* was observed in medial sections of the *Ezh2* cKO animals (**Figure 8A**, middle panel, arrow) and levels of *Wnt1* were significantly lower in *En1cre/+; Ezh2* L/L animals compared to wildtypes (n=3, P< 0.05, one-tailed) (**Figure 8B**). Another important determinant for the IsO is *Otx2*. As described above, *Otx2* plays a role in positioning the IsO, as the caudal limit of the *Otx2* expression domain marks the location of the IsO (Broccoli et al., 1999a; Wurst and Bally-Cuif, 2001). In contrast to *Fgf8* and *Wnt1, Otx2* expression was found to be expanded caudally into the hindbrain of *Ezh2* cKO embryos (**Figure 8AC**, lower panels, arrowheads). Normally *Otx2* expression is not found caudal to *Fgf8* expression, however in medial sections of *En1cre/+; Ezh2* L/L embryos we could observe expression of *Otx2* caudal to the *Fgf8* expression domain (**Figure 8C**, arrowheads). Even though an expansion of the expression domain of *Otx2* is visible, the overall levels of *Otx2* were found to be reduced in the *Ezh2* mutant (n=3, P<0.01, one-tailed) (**Figure 8B**). To further substantiate the caudal *Otx2* expression, we also performed *in situ* hybridization for *Otx2* on E14.5 *Wnt1Cre/+; Ezh2* L/L embryos (**Supplemental figure 2**). Using the *Th* expression domain from **Figure 6B** as a reference, a patch of *Otx2* expression could be observed caudal to the midbrain in medial sections of *Wnt1Cre* driven *Ezh2* cKOs, which was not present in wildtype littermates (**Supplemental figure 2**, arrowhead). Together these data indicate that *Ezh2* plays a role in suppressing *Otx2* expression in the R1 region, next to influencing expression levels of *Fgf8* and *Wnt1*.

## Discussion

A critical early event in the generation of different neuronal subtypes is the regionalization of the neural tube. This process in the mid-hindbrain region is dependent on the signaling factors produced by the IsO (Crossley et al., 1996; Wurst and Bally-Cuif, 2001). The IsO is formed at the juxtaposition of *Otx2/Gbx2* expression around ~E7-8 (Broccoli et al., 1999b; Joyner et al., 2000), and depends on several transcription factors for its proper functioning and maintenance (Adams et al., 2000; Canning et al., 2007; Guo et al., 2006; Li et al., 2002). In the current study we show that next to transcription factors, transcriptional inhibition via PRC2 and H3K27me3 also play a role in maintaining the IsO and determining the fate of cells in the mid- hindbrain region. The loss of *Ezh2* and consequently H3K27me3 affected the molecular coding of the anterior R1. DA-related genes were found ectopically in the ventral anterior hindbrain and a caudal expansion of the TH positive population was observed. The emergence of DA neurons in the Isthmic area was at the expanse of 5-HT neurons and *Nkx6.1* expression in the trochlear motor neurons located directly caudal to the IsO.

The deletion of *Ezh2* mimics the phenotype of the *En1* mutant, in which a disorganized IsO causes a posterior expansion of the midbrain a the expanse of the anterior ventral R1 region (Kouwenhoven et al., 2016). Even though both proteins might mechanistically function via the regulation of *Otx2*, no direct relationship between the two factors was found. The expression of *En1* is not affected in *En1cre/ +; Ezh2* L/L embryos and the results were verified in a second independent model, *Wnt1Cre/Ezh2.* In addition, no global changes were observed in H3K27me3 levels in *En1* mutants compared to wildtypes, suggesting that general PRC2 functioning is not impaired. However, the targeting of the PRC2 complex might be altered in *En1* mutants, leading to local changes in H3K27me3 levels, which are not visualized by overall immunohistochemistry for H3K27me3. Furthermore, analysis of the TH+ domain in *En1/Ezh2* double mutants does imply that EZH2 and EN1 have a phenotypic relationship as the expansion of the TH+ region is aggravated in animals depleted of both *En1* and *Ezh2*.

The induction of DA neurons in R1 of *Ezh2* cKOs can be explained by the presence of *Otx2* in the hindbrain (Brodski et al., 2003; Sherf et al., 2015). At E12.5 we observed *Otx2* expression caudal to the midbrain in *En1cre/+; Ezh2* L/L embryos and *Wnt1Cre* driven *Ezh2* cKOs also displayed a caudal expansion of the *Otx2* domain. It has been shown that repression of *Otx2* in the hindbrain initially depends on GBX2, however after E8.5 a GBX2-independent mechanism can repress *Otx2* in R1 (Li et al., 2002). At E12.5 the *Fgf8* expression domain marks the caudal limit of the *Otx2* expression domain and a study in which a FGF8-bead was implanted in p2, showed a regional repression of *Otx2*, suggesting that FGF8 can compensate for the loss of GBX2 in repressing *Otx2* after E8.5 (Li et al., 2002). However, the introduction of FGF8 into this region also led to the induction of several other IsO related genes, including *En1*, which might also contribute to the repression of *Otx2* (Shamim et al., 1999). Although the expansion of *Otx2* into the hindbrain might additionally be a consequence of alterations in IsO related genes, we argue that the direct loss of H3K27me3 on the promoter of *Otx2* may cause the observed reactivation of the *Otx2* gene in our model. *Otx2* has already been identified as a direct target of PRC2 in embryonic stem cells (Boyer et al., 2006; Lee et al., 2006; Pasini et al., 2007). Loss of a functional PRC2 complex led to the loss of H3K27me3 on the promoter of *Otx2* and an increase in its expression level (Pasini et al., 2007, Supplemental figure 1). In addition, H3K27me3 is found on the promoter of *Otx2* in cell types that do not express *Otx2* (**Supplemental figure 3**), suggesting that the silencing of *Otx2* might be H3K27me3-mediated (Mikkelsen et al., 2007; Mohn et al., 2008, https://genome.ucsc.edu/index.html; mouse NCBI37/mm9). Interestingly, in contrast to previous studies the altered expression of *Otx2* was not mimicked by *Wnt1* in *En1cre/+; Ezh2* L/L animals (Broccoli et al., 1999a; Kouwenhoven et al., 2016)(Broccoli et al., 1999b; Kouwenhoven et al., 2016). The levels of *Wnt1* were down-regulated in *En1cre* driven *Ezh2* mutants and reduced spatial expression was observed in medial sections of E12.5 *En1cre/+; Ezh2* L/L embryos. In addition, reduced canonical Wnt signaling was found in *Wnt1Cre/+; Ezh2* L/L embryos (Zemke et al., 2015), suggesting that the sole presence of *Otx2* in the hindbrain of *Ezh2* cKOs is enough to fate switch the hindbrain R1 region towards a midbrain phenotype.

Taken together, our data show that next to transcription factors, transcriptional regulation via epigenetics also plays an important part in the development of the Isthmic area. Deletion of *Ezh2* leads to *Otx2* expression in the hindbrain and the induction of DA neurons at the expense of 5-HT positive and Nkx6.1 positive neurons. The observed phenotype mimics that of *En1* mutant animals and even though *Ezh2* and *En1* do not appear to have a direct regulatory relationship, we theorize that EN1 might be required for the targeting of EZH2 and PRC2 to the *Otx2* locus.

## Acknowledgments

We would like to thank Dr. Frederic Zilbermann of the Friedrich Miescher Institute for his generous gift of the *Ezh2*-floxed mouse line and Dr. Sandra Blaess for kindly providing us with the *En1Cre; R26RYFP* mouse strain. In addition we would like to thank Prof. Dr. Filippo Rijli for sending us E14.5 frozen and PFA/Sucrose *Wnt1Cre; Ezh2* embryos.

---

**Supplemental figure 1:**
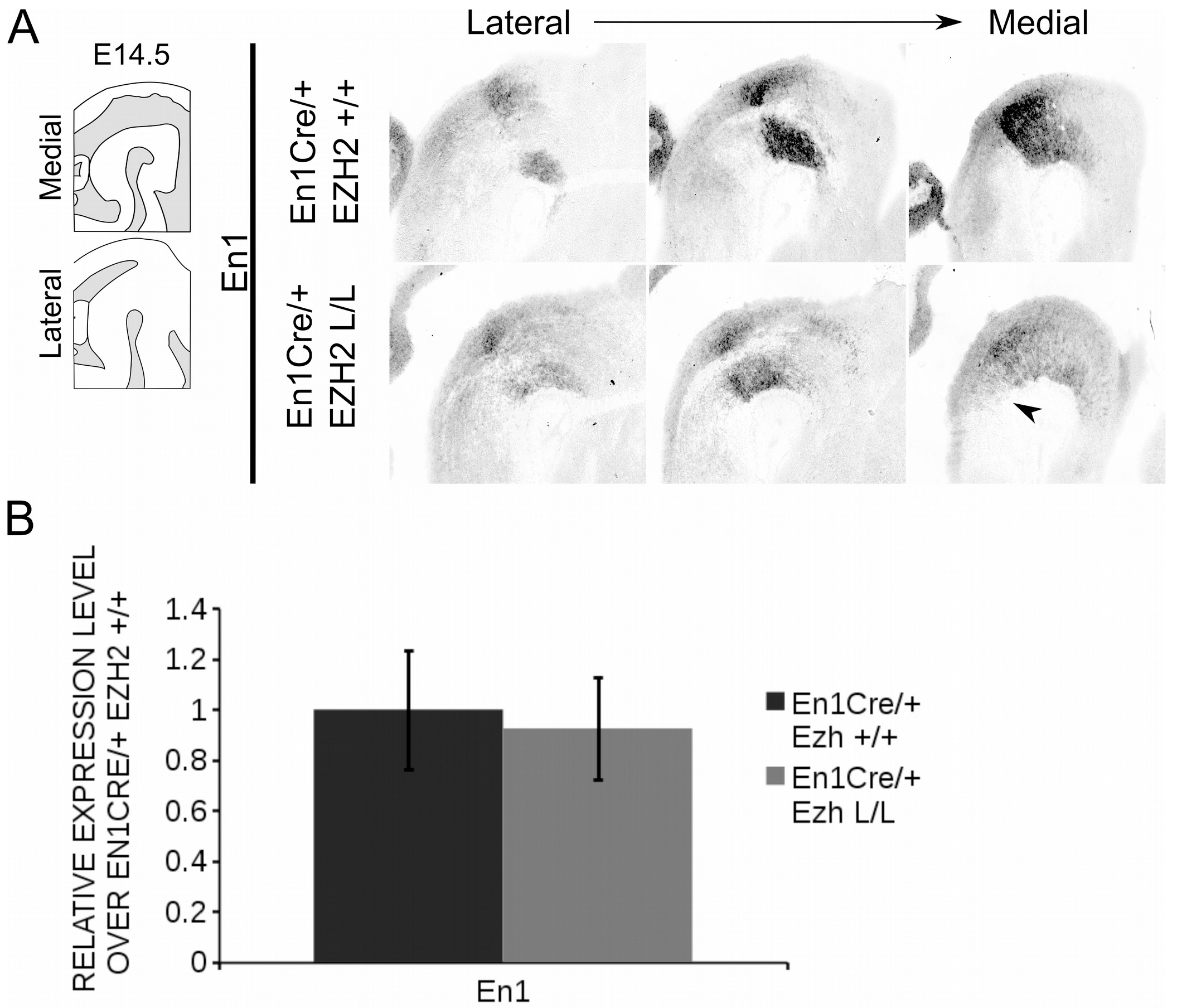
The loss of *Ezh2* does not influence *En1* expression. The expression of *En1* was analyzed with *in situ* hybridization and qPCR at E14.5 (A, B). (A) *En1* shows a similar expression pattern between the *En1cre/+; Ezh2 +/+* and the *En1cre/+; Ezh2* L/L animals. However ectopic expression of *En1* might be present in the ventral region of R1 (arrowhead).(B) The levels of *En1* are not significantly changed between wildtypes and *Ezh2* cKO animals at E14.5.

**Supplemental figure 2:**
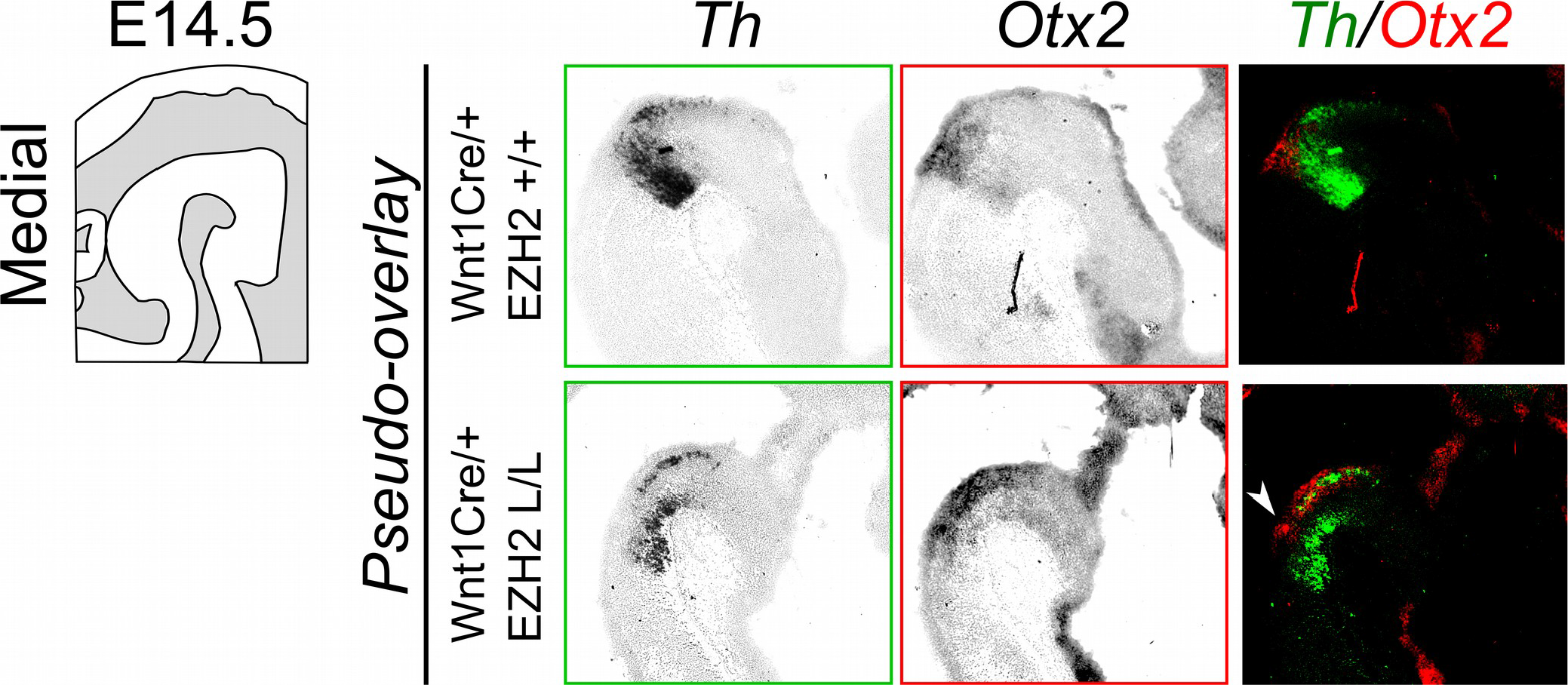
*Otx2* is found caudal to the midbrain in *Wnt1Cre/+; Ezh2* L/L embryos. *Otx2* expression was assessed by means of *in situ* hybridization using *Th* expression as a reference for the location. A pseudo-overlay of adjacent slides of demonstrates the presence of *Otx2* expression dorsal of the ectopic *Th* expression in medial sections of *Wnt1Cre* driven *Ezh2* cKOs (arrowhead).

**Supplemental figure 3:**
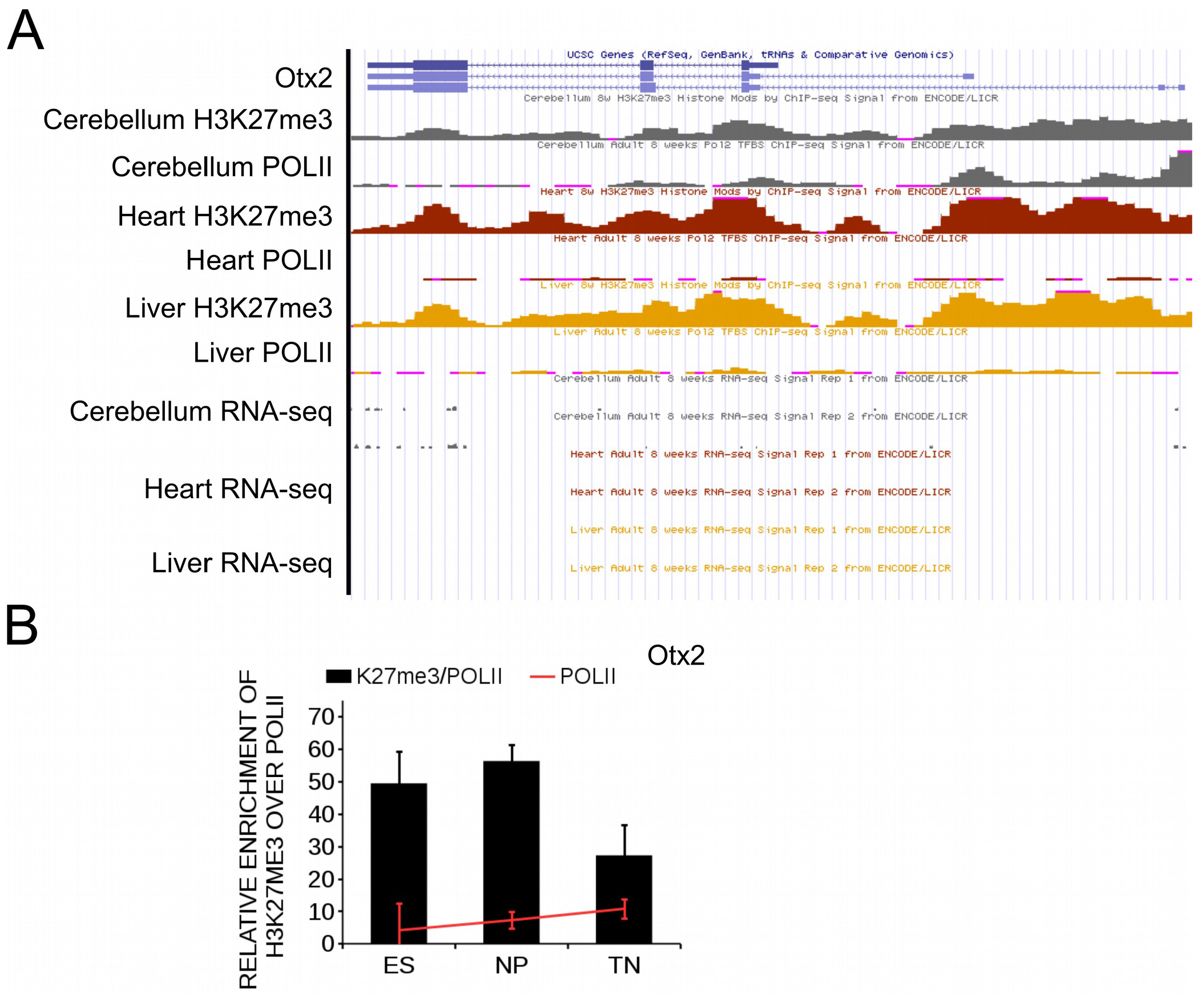
H3K27me3 is present on the promoter of *Otx2* in tissue in which *Otx2* is not expressed. (A) Visualization of ChIP-sequencing peaks of H3K27me3 and RNA-polymerase II (POLII) at the *Otx2* promoter obtained from https://genome.ucsc.edu/index.html using the mouse NCBI37/mm9 assembly. In tissue where almost no RNA-sequencing reads are detected for *Otx2*, a high peak for H3K27me3 is observed, while the peak for POLII is low. (B) Analysis of ChIP-sequencing data from (Mohn et al., 2008). The Otx2 promoter shows a high enrichment for H3K27me3 in Embryonic stem cells (ES), Neuronal progenitors (NP) and Terminal pyramidal glutamatergic neurons (TN), while enrichment for POLII is low, indicating that *Otx2* is not activated in these cells.

## References

Adams, K.A., Maida, J.M., Golden, J.A., and Riddle, R.D. (2000). The transcription factor Lmxlb maintains Wntl expression within the isthmic organizer. Development 127, 1857–1867.

Alonso, A., Merchán, P., Sandoval, J.E., Sánchez-Arrones, L., Garcia-Cazorla, A., Artuch, R., Ferrán, J.L., Martínez-de-la-Torre, M., and Puelles, L. (2013). Development of the serotonergic cells in murine raphe nuclei and their relations with rhombomeric domains. Brain Struct. Funct. 218, 1229–1277.

Andersson, E., Tryggvason, U., Deng, Q., Friling, S., Alekseenko, Z., Robert, B., Perlmann, T., and Ericson, J. (2006). Identification of Intrinsic Determinants of Midbrain Dopamine Neurons. Cell 124, 393–405.

Arenas, E., Denham, M., and Villaescusa, J.C. (2015). How to make a midbrain dopaminergic neuron. Development 142, 1918–1936.

Bayer, S.A., Wills, K.V., Triarhou, L.C., and Ghetti, B. (1995). Time of neuron origin and gradients of neurogenesis in midbrain dopaminergic neurons in the mouse. Exp. Brain Res. 105, 191–199.

Bernstein, B.E., Mikkelsen, T.S., Xie, X., Kamal, M., Huebert, D.J., Cuff, J., Fry, B., Meissner, A., Wernig, M., Plath, K., et al. (2006). A Bivalent Chromatin Structure Marks Key Developmental Genes in Embryonic Stem Cells. Cell 125, 315–326.

Boyer, L.A., Plath, K., Zeitlinger, J., Brambrink, T., Medeiros, L.A., Lee, T.I., Levine, S.S., Wernig, M., Tajonar, A., Ray, M.K., et al. (2006). Polycomb complexes repress developmental regulators in murine embryonic stem cells. Nature 441, 349–353.

Broccoli, V., Boncinelli, E., and Wurst, W. (1999a). The caudal limit of Otx2 expression positions the isthmic organizer. Nature 401, 164–168.

Broccoli, V., Boncinelli, E., and Wurst, W. (1999b). The caudal limit of Otx2 expression positions the isthmic organizer. Nature 401, 164–168.

Brodski, C., Weisenhorn, D.M.V., Signore, M., Sillaber, I., Oesterheld, M., Broccoli, V., Acampora, D., Simeone, A., and Wurst, W. (2003). Location and Size of Dopaminergic and Serotonergic Cell Populations Are Controlled by the Position of the Midbrain-Hindbrain Organizer. J. Neurosci. 23, 4199–4207.

Canning, C.A., Lee, L., Irving, C., Mason, I., and Jones, C.M. (2007). Sustained interactive Wnt and FGF signaling is required to maintain isthmic identity. Dev. Biol. 305, 276–286.

Cao, R., and Zhang, Y. (2004a). SUZ12 Is Required for Both the Histone Methyltransferase Activity and the Silencing Function of the EED-EZH2 Complex. Mol. Cell 15, 57–67.

Cao, R., and Zhang, Y. (2004b). The functions of E(Z)/EZH2-mediated methylation of lysine 27 in histone H3. Curr. Opin. Genet. Dev. 14, 155–164.

Cedar, H., and Bergman, Y. (2009). Linking DNA methylation and histone modification: patterns and paradigms. Nat. Rev. Genet. 10, 295.

Chandrasekhar, A. (2004). Turning Heads: Development of Vertebrate Branchiomotor Neurons. Dev. Dyn. Off. Publ. Am. Assoc. Anat. 229, 143–161.

Corley, M., and Kroll, K.L. (2015). The Roles and Regulation of Polycomb Complexes in Neural Development. Cell Tissue Res. 359, 65–85.

Crossley, P.H., Martinez, S., and Martin, G.R. (1996). Midbrain development induced by FGF8 in the chick embryo. Nature 380, 66–68.

Danielian, P.S., and McMahon, A.P. (1996). Engrailed-1 as a target of the Wnt-1 signalling pathway in vertebrate midbrain development. Nature 383, 332–334.

Deng, Q., Andersson, E., Hedlund, E., Alekseenko, Z., Coppola, E., Panman, L., Millonig, J.H., Brunet, J.-F., Ericson, J., and Perlmann, T. (2011). Specific and integrated roles of Lmx1a, Lmx1b and Phox2a in ventral midbrain development. Development 138, 3399–3408.

Faust, C., Lawson, K.A., Schork, N.J., Thiel, B., and Magnuson, T. (1998). The Polycomb-group gene eed is required for normal morphogenetic movements during gastrulation in the mouse embryo. Development 125, 4495–4506.

Ferrari, K.J., Scelfo, A., Jammula, S., Cuomo, A., Barozzi, I., Stützer, A., Fischle, W., Bonaldi, T., and Pasini, D. (2014). Polycomb-Dependent H3K27me1 and H3K27me2 Regulate Active Transcription and Enhancer Fidelity. Mol. Cell 53, 49–62.

Fox, S.R., and Deneris, E.S. (2012). Engrailed is required in maturing serotonin neurons to regulate the cytoarchitecture and survival of the dorsal raphe nucleus. J. Neurosci. Off. J. Soc. Neurosci. 32, 7832–7842.

Grima, B., Lamouroux, A., Blanot, F., Biguet, N.F., and Mallet, J. (1985). Complete coding sequence of rat tyrosine hydroxylase mRNA. Proc. Natl. Acad. Sci. U. S. A. 82, 617–621.

Guo, C., Qiu, H.-Y., Huang, Y., Chen, H., Yang, R.-Q., Chen, S.-D., Johnson, R.L., Chen, Z.-F., and Ding, Y.-Q. (2006). <em>Lmx1b</em> is essential for <em>Fgf8</em> and <em>Wnt1</em> expression in the isthmic organizer during tectum and cerebellum development in mice. Development 134, 317.

Heesbeen, H.J. van, Mesman, S., Veenvliet, J.V., and Smidt, M.P. (2013). Epigenetic mechanisms in the development and maintenance of dopaminergic neurons. Development 140, 1159–1169.

Hirabayashi, Y., Suzki, N., Tsuboi, M., Endo, T.A., Toyoda, T., Shinga, J., Koseki, H., Vidal, M., and Gotoh, Y. (2009). Polycomb Limits the Neurogenic Competence of Neural Precursor Cells to Promote Astrogenic Fate Transition. Neuron 63, 600–613.

Hoekstra, E.J., von Oerthel, L., van der Heide, L.P., Kouwenhoven, W.M., Veenvliet, J.V., Wever, I., Jin, Y.-R., Yoon, J.K., van der Linden, A.J.A., Holstege, F.C.P., et al. (2013). Lmx1a Encodes a Rostral Set of Mesodiencephalic Dopaminergic Neurons Marked by the Wnt/B-Catenin Signaling Activator R-spondin 2. PLoS ONE 8.

Hwang, D.-Y., Ardayfio, P., Kang, U.J., Semina, E.V., and Kim, K.-S. (2003). Selective loss of dopaminergic neurons in the substantia nigra of Pitx3-deficient aphakia mice. Mol. Brain Res. 114, 123–131.

Iversen, L.L. (2010). Dopamine Handbook (Oxford University Press).

Jacobs, F.M.J., Smits, S.M., Noorlander, C.W., Oerthel, L. von, Linden, A.J.A. van der, Burbach, J.P.H., and Smidt, M.P. (2007). Retinoic acid counteracts developmental defects in the substantia nigra caused by Pitx3 deficiency. Development 134, 2673–2684.

Jacobs, F.M.J., Veenvliet, J.V., Almirza, W.H., Hoekstra, E.J., Oerthel, L.von, Linden, A.J.A. van der, Neijts, R., Koerkamp, M.G., Leenen, D.van, Holstege, F.C.P., et al. (2011). Retinoic acid-dependent and -independent gene-regulatory pathways of Pitx3 in meso-diencephalic dopaminergic neurons. Development 138, 5213–5222.

Jankovic, J., Chen, S., and Le, W.D. (2005). The role of Nurr1 in the development of dopaminergic neurons and Parkinson’s disease. Prog. Neurobiol. 77, 128–138.

Joyner, A.L., Liu, A., and Millet, S. (2000). Otx2, Gbx2 and Fgf8 interact to position and maintain a mid-hindbrain organizer. Curr. Opin. Cell Biol. 12, 736–741.

Kawano, H., Ohyama, K., Kawamura, K., and Nagatsu, I. (1995). Migration of dopaminergic neurons in the embryonic mesencephalon of mice. Brain Res. Dev. Brain Res. 86, 101–113.

Kimmel, R.A., Turnbull, D.H., Blanquet, V., Wurst, W., Loomis, C.A., and Joyner, A.L. (2000). Two lineage boundaries coordinate vertebrate apical ectodermal ridge formation. Genes Dev. 14, 1377–1389.

Kouwenhoven, W.M., Veenvliet, J.V., Hooft, J.A.van, Heide, L.P. van der, and Smidt, M.P. (2016). Engrailed 1 shapes the dopaminergic and serotonergic landscape through proper isthmic organizer maintenance and function. Biol. Open 5, 279–288.

Kratochwil, C.F., Maheshwari, U., and Rijli, F.M. (2017). The Long Journey of Pontine Nuclei Neurons: From Rhombic Lip to Cortico-Ponto-Cerebellar Circuitry. Front. Neural Circuits 11.

Lee, T.I., Jenner, R.G., Boyer, L.A., Guenther, M.G., Levine, S.S., Kumar, R.M., Chevalier, B., Johnstone, S.E., Cole, M.F., Isono, K., et al. (2006). Control of Developmental Regulators by Polycomb in Human Embryonic Stem Cells. Cell 125, 301–313.

Li, J.Y.H., Lao, Z., and Joyner, A.L. (2002). Changing Requirements for Gbx2 in Development of the Cerebellum and Maintenance of the Mid/Hindbrain Organizer. Neuron 36, 31–43.

Margueron, R., Li, G., Sarma, K., Blais, A., Zavadil, J., Woodcock, C.L., Dynlacht, B.D., and Reinberg, D. (2008). Ezh1 and Ezh2 Maintain Repressive Chromatin through Different Mechanisms. Mol. Cell 32, 503–518.

Martinez, S., Crossley, P.H., Cobos, I., Rubenstein, J.L., and Martin, G.R. (1999). FGF8 induces formation of an ectopic isthmic organizer and isthmocerebellar development via a repressive effect on Otx2 expression. Development 126, 1189–1200.

Maxwell, S.L., Ho, H.-Y., Kuehner, E., Zhao, S., and Li, M. (2005). Pitx3 regulates tyrosine hydroxylase expression in the substantia nigra and identifies a subgroup of mesencephalic dopaminergic progenitor neurons during mouse development. Dev. Biol. 282, 467–479.

McMahon, A.P., Joyner, A.L., Bradley, A., and McMahon, J.A. (1992). The midbrain-hindbrain phenotype of Wnt-1–Wnt-1–mice results from stepwise deletion of engrailed-expressing cells by 9.5 days postcoitum. Cell 69, 581–595.

Mesman, S., von Oerthel, L., and Smidt, M.P. (2014). Mesodiencephalic Dopaminergic Neuronal Differentiation Does Not Involve GLI2A-Mediated SHH-Signaling and Is under the Direct Influence of Canonical WNT Signaling. PLoS ONE 9.

Mikkelsen, T.S., Ku, M., Jaffe, D.B., Issac, B., Lieberman, E., Giannoukos, G., Alvarez, P., Brockman, W., Kim, T.-K., Koche, R.P., et al. (2007). Genome-wide maps of chromatin state in pluripotent and lineage-committed cells. Nature 448, 553–560.

Mohn, F., Weber, M., Rebhan, M., Roloff, T.C., Richter, J., Stadler, M.B., Bibel, M., and Schübeler, D. (2008). Lineage-Specific Polycomb Targets and De Novo DNA Methylation Define Restriction and Potential of Neuronal Progenitors. Mol. Cell 30, 755–766.

O’Carroll, D., Erhardt, S., Pagani, M., Barton, S.C., Surani, M.A., and Jenuwein, T. (2001). The Polycomb-Group GeneEzh2 Is Required for Early Mouse Development. Mol. Cell. Biol. 21, 4330–4336.

Omodei, D., Acampora, D., Mancuso, P., Prakash, N., Giovannantonio, L.G.D., Wurst, W., and Simeone, A. (2008). Anterior-posterior graded response to Otx2 controls proliferation and differentiation of dopaminergic progenitors in the ventral mesencephalon. Development 135, 3459–3470.

Ono, Y., Nakatani, T., Sakamoto, Y., Mizuhara, E., Minaki, Y., Kumai, M., Hamaguchi, A., Nishimura, M., Inoue, Y., Hayashi, H., et al. (2007). Differences in neurogenic potential in floor plate cells along an anteroposterior location: midbrain dopaminergic neurons originate from mesencephalic floor plate cells. Development 134, 3213–3225.

Pasini, D., Bracken, A.P., Hansen, J.B., Capillo, M., and Helin, K. (2007). The Polycomb Group Protein Suz12 Is Required for Embryonic Stem Cell Differentiation. Mol. Cell. Biol. 27, 3769–3779.

Pereira, J.D., Sansom, S.N., Smith, J., Dobenecker, M.-W., Tarakhovsky, A., and Livesey, F.J. (2010). Ezh2, the histone methyltransferase of PRC2, regulates the balance between self-renewal and differentiation in the cerebral cortex. Proc. Natl. Acad. Sci. 107, 15957–15962.

Placzek, M., and Briscoe, J. (2005). The floor plate: multiple cells, multiple signals. Nat. Rev. Neurosci. 6, 230–240.

Podobinska, M., Szablowska-Gadomska, I., Augustyniak, J., Sandvig, I., Sandvig, A., and Buzanska, L. (2017). Epigenetic Modulation of Stem Cells in Neurodevelopment: The Role of Methylation and Acetylation. Front. Cell. Neurosci. 11.

Prakash, N., Puelles, E., Freude, K., Trümbach, D., Omodei, D., Salvio, M.D., Sussel, L., Ericson, J., Sander, M., Simeone, A., et al. (2009). Nkx6-1 controls the identity and fate of red nucleus and oculomotor neurons in the mouse midbrain. Development 136, 2545–2555.

Saucedo-Cardenas, O., Quintana-Hau, J.D., Le, W.-D., Smidt, M.P., Cox, J.J., Mayo, F.D., Burbach, J.P.H., and Conneely, O.M. (1998). Nurr1 is essential for the induction of the dopaminergic phenotype and the survival of ventral mesencephalic late dopaminergic precursor neurons. Proc. Natl. Acad. Sci. 95, 4013–4018.

Shamim, H., Mahmood, R., Logan, C., Doherty, P., Lumsden, A., and Mason, I. (1999). Sequential roles for Fgf4, En1 and Fgf8 in specification and regionalisation of the midbrain. Development 126, 945–959.

Shen, X., Liu, Y., Hsu, Y.-J., Fujiwara, Y., Kim, J., Mao, X., Yuan, G.-C., and Orkin, S.H. (2008). EZH1 Mediates Methylation on Histone H3 Lysine 27 and Complements EZH2 in Maintaining Stem Cell Identity and Executing Pluripotency. Mol. Cell 32, 491–502.

Sherf, O., Nashelsky Zolotov, L., Liser, K., Tilleman, H., Jovanovic, V.M., Zega, K., Jukic, M.M., and Brodski, C. (2015). Otx2 Requires Lmx1b to Control the Development of Mesodiencephalic Dopaminergic Neurons. PLoS ONE 10.

Shults, C.W., Hashimoto, R., Brady, R.M., and Gage, F.H. (1990). Dopaminergic cells align along radial glia in the developing mesencephalon of the rat. Neuroscience 38, 427–436.

Smidt, M.P., van Schaick, H.S.A., Lanctôt, C., Tremblay, J.J., Cox, J.J., van der Kleij, A.A.M., Wolterink, G., Drouin, J., and Burbach, J.P.H. (1997). A homeodomain gene Ptx3 has highly restricted brain expression in mesencephalic dopaminergic neurons. Proc. Natl. Acad. Sci. U. S. A. 94, 13305–13310.

Smidt, M.P., Asbreuk, C.H.J., Cox, J.J., Chen, H., Johnson, R.L., and Burbach, J.P.H. (2000). A second independent pathway for development of mesencephalic dopaminergic neurons requires Lmx1b. Nat. Neurosci. 3, 337–341.

Smidt, M.P., Smits, S.M., Bouwmeester, H., Hamers, F.P.T., Linden, A.J.A. van der, Hellemons, A.J.C.G.M., Graw, J., and Burbach, J.P.H. (2004). Early developmental failure of substantia nigra dopamine neurons in mice lacking the homeodomain gene Pitx3. Development 131, 1145–1155.

Smits, S.M., Ponnio, T., Conneely, O.M., Burbach, J.P.H., and Smidt, M.P. (2003). Involvement of Nurr1 in specifying the neurotransmitter identity of ventral midbrain dopaminergic neurons. Eur. J. Neurosci. 18, 1731–1738.

Srinivas, S., Watanabe, T., Lin, C.S., William, C.M., Tanabe, Y., Jessell, T.M., and Costantini, F. (2001). Cre reporter strains produced by targeted insertion of EYFP and ECFP into the ROSA26 locus. BMC Dev. Biol. 1, 4.

Sunmonu, N.A., Chen, L., and Li, J.Y.H. (2009). Misexpression of Gbx2 throughout the mesencephalon by a conditional gain-of-function transgene leads to deletion of the midbrain and cerebellum in mice. Genes. N. Y. N 2000 47, 667–673.

Veenvliet, J.V., Santos, M.T.M.A.dos, Kouwenhoven, W.M., Oerthel, L.von, Lim, J.L., Linden, A.J.A. van der, Koerkamp, M.J.A.G., Holstege, F.C.P., and Smidt, M.P. (2013). Specification of dopaminergic subsets involves interplay of En1 and Pitx3. Development 140, 4116–4116.

Wurst, W., and Bally-Cuif, L. (2001). Neural plate patterning: Upstream and downstream of the isthmic organizer. Nat. Rev. Neurosci. 2, 99–108.

Yan, C.H., Levesque, M., Claxton, S., Johnson, R.L., and Ang, S.-L. (2011). Lmx1a and Lmx1b Function Cooperatively to Regulate Proliferation, Specification, and Differentiation of Midbrain Dopaminergic Progenitors. J. Neurosci. 31, 12413–12425.

Zemke, M., Draganova, K., Klug, A., Schöler, A., Zurkirchen, L., Gay, M.H.-P., Cheng, P., Koseki, H., Valenta, T., Schübeler, D., et al. (2015). Loss of Ezh2 promotes a midbrain-to-forebrain identity switch by direct gene derepression and Wnt-dependent regulation. BMC Biol. 13, 103.

